# Automated in vivo tracking of cortical oligodendrocytes

**DOI:** 10.1101/2021.02.12.430879

**Authors:** Yu Kang T. Xu, Cody L. Call, Jeremias Sulam, Dwight E. Bergles

## Abstract

Oligodendrocytes exert a profound influence on neural circuits by accelerating axon potential conduction, altering excitability and providing metabolic support. As oligodendrogenesis continues in the adult brain and is essential for myelin repair, uncovering the factors that control their dynamics is necessary to understand the consequences of adaptive myelination and develop new strategies to enhance remyelination in diseases such as multiple sclerosis. Unfortunately, few methods exist for analysis of oligodendrocyte dynamics, and even fewer are suitable for *in vivo* investigation. Here, we describe the development of a fully automated cell tracking pipeline using convolutional neural networks (*Oligo-Track*) that provides rapid volumetric segmentation and tracking of thousands of cells over weeks *in vivo*. This system reliably replicated human analysis, outperformed traditional analytic approaches, and extracted injury and repair dynamics at multiple cortical depths, establishing that oligodendrogenesis after cuprizone-mediated demyelination is suppressed in deeper cortical layers. Volumetric data provided by this analysis revealed that oligodendrocyte soma size progressively decreases after their generation, and declines further prior to death, providing a means to predict cell age and eventual cell death from individual time points. This new CNN-based analysis pipeline offers a rapid, robust method to quantitatively analyze oligodendrocyte dynamics *in vivo*, which will aid in understanding how changes in these myelinating cells influence circuit function and recovery from injury and disease.

## INTRODUCTION

Advances in genetically encoded fluorescent indicators, CRISPR-mediated gene editing and multiphoton microscopy provide unprecedented opportunities for studying cellular dynamics at single-cell resolution in the brains of living animals. While these approaches hold the potential for profound discoveries about brain function, they also come with a host of quantitative challenges. In particular, living brain tissue is unstable; tissue warping disrupts image quality and uneven refractive indices increase noise and produce anisotropic distortions during longitudinal image acquisition (Lecoq et al., 2019). Moreover, large multi-dimensional datasets are cumbersome to quantify, and often require specialized software for 4D visualization and manual curation (Pidhorskyi et al., 2018). As imaging tools become more advanced and enable researchers to delve deeper into the brain *in vivo* (Horton et al., 2013), the challenges associated with quantification of enormous datasets become more acute. Further advances depend critically on the availability of robust analysis platforms to rapidly extract multi-dimensional observations about cellular dynamics.

Developing rigorous analysis tools for *in vivo* investigation of oligodendrocytes is particularly important. Oligodendrocytes enhance the speed of action potential conduction by ensheathing neuronal axons with concentric wraps of membrane, support neuronal metabolism and control neuronal excitability (Simons and Nave, 2016; Larson et al., 2018). While the population of neurons in the brain remains relatively stable throughout life (Bhardwaj et al., 2006; Ming and Song, 2011), new oligodendrocytes are generated in the adult CNS, allowing for dynamic alteration of myelin patterns in both healthy and pathological conditions (El Waly et al., 2014). This dynamism highlights the need for automated, longitudinal tracking tools to quantify the location, timing and extent of myelin plasticity within defined circuits in response to particular behavioral paradigms, as well as the regeneration of oligodendrocytes after demyelination (Bergles and Richardson, 2015). In this study, we sought to develop fully automated methodologies to overcome the analytic challenges associated with longitudinal tracking of oligodendrocytes *in vivo*.

Currently, most available cell tracking algorithms are designed for *in vitro* analysis and are not readily adaptable to *in vivo* conditions (Van Valen et al., 2016; Zhong et al., 2016; Nketia et al., 2017; Lugagne et al., 2020; Wang et al., 2020). The few *in vivo* tracking algorithms that exist are modality specific and cannot be readily adapted to our fluorescent longitudinal datasets (Acton et al., 2002; Nguyen et al., 2011). The closest *in vivo* tools that can be applied to oligodendrocyte datasets are those developed for analyzing calcium imaging (Pachitariu et al., 2017; Giovannucci et al., 2019). However, calcium imaging tools normally work best with high-frame rate videos taken over seconds, rather than image volumes collected on a weekly basis that often experience large-scale tissue warping between imaging sessions. To resolve this longitudinal volumetric tracking challenge, we opted to use convolutional neural networks (CNN), which are known to find accurate efficient solutions to high-dimensional problems. Convolutional kernels allow CNNs to adaptively assess local features and global spatial relationships to make tracking decisions that are more perceptual, or human-like. Moreover, additional techniques such as transfer learning can help trained models generalize to entirely new imaging challenges with minimal new training data (Zhuang et al., 2020), extending their use to other contexts.

Here, we describe the development of *Oligo-Track*, a fast and reliable cell tracker for *in vivo* semantic segmentation of oligodendrocyte dynamics across cortical layers in longitudinal imaging experiments. We validated our algorithm using the cuprizone model of demyelination *in vivo* and show that *Oligo-Track* outperforms traditional analytic approaches in extracting dynamics of oligodendrogenesis at greater depths than previously available with manual annotation. Moreover, this approach generated volumetric segmentations of tracked cells that were inaccessible to human analysis, due to the considerable time investment required for manual volumetric tracing. This volumetric data revealed that oligodendrocyte soma size varies predictably with age and proximity to death, allowing additional information about the timing of oligodendrogenesis and cell death to be extracted from fixed timepoint imaging experiments.

## MATERIALS and METHODS

### Animal care and use

Female and male adult mice were used for experiments and randomly assigned to experimental groups. All mice were healthy and did not display any overt behavioral phenotypes, and no animals were excluded from the analysis. Generation and genotyping of BAC transgenic lines from *Mobp-EGFP* (GENSAT) have been previously described (Hughes et al., 2018). Mice were maintained on a 12 hr light/dark cycle, housed in groups no larger than 5, and food and water were provided ad libitum (except during cuprizone-administration, see below). All animal experiments were performed in strict accordance with protocols approved by the Animal Care and Use Committee at Johns Hopkins University.

### Cranial windows

Cranial windows were prepared as previously described (Holtmaat et al., 2012; Hughes et al., 2018; Orthmann-Murphy et al., 2020). Mice aged 7 to 10 weeks were deeply anesthetized with isoflurane (5% with 1 L/min O_2_ induction; 1.5–2% with 0.5 L/min maintenance), the head shaved, and the scalp removed to expose the skull. The skull was cleaned and dried and a position over somatosensory cortex (−1.5 mm posterior and 3.5 mm lateral from bregma) was marked for drilling. A custom aluminum headplate with a central hole was cemented onto the skull (C and B Metabond) and fixed in place with custom clamping headbars. A 2 mm × 2 mm square or 3 mm × 3 mm circle of skull was removed using a high-speed dental drill. A coverslip (VWR, No. 1) the size of the craniotomy was put in its place and sealed with cyanoacrylate glue (Vetbond and Krazy glue).

### *In vivo* two photon microscopy

*In vivo* imaging was performed as previously described (Orthmann-Murphy et al., 2020). After two to three weeks of recovery from cranial window surgery, baseline images of the cortex were acquired with two photon microscopy on a Zeiss LSM 710 microscope (average power at sample < 30 mW). Image stacks were 425 μm × 425 μm × 550 μm or 850 μm × 850 μm × 550 μm (1024 × 1024 pixels; corresponding to layers I – IV), relative to the pia. Mice were subsequently imaged weekly for up to 12 weeks.

### Cuprizone treatment

Directly following baseline two photon image acquisition, mice were switched from regular diet to a diet consisting of milled, irradiated 18% protein chow (Teklad Global) supplemented with 0.2% w/w bis(cyclohexanone) oxaldihydrazone (“cuprizone,” Sigma). Control mice received only the milled chow. After three weeks, mice returned to regular pellet diet for the duration of the recovery period (Orthmann-Murphy et al., 2020).

### Analytic pipeline overview

Timeseries acquired from our two-photon imaging setup were first registered using ImageJ’s correct 3D drift plugin (Schindelin et al., 2012; Parslow et al., 2014), which accounted for major alignment shifts from week to week. Registered timeseries were then analyzed crop-by-crop using our segmentation CNN (Seg-CNN) which identified cell somas on a voxel-wise basis. These cell somas were then extracted as individual seeds for our tracking CNN (Track-CNN) that identified the location of each seeded cell soma on a subsequent time point. In parallel, we also developed a cell tracking method based on traditional imaging informatics approaches that used the structural similarity index (SSIM) (Zhou Wang et al., 2004) and local tissue movement calculations to track cells. This heuristic model was used as a baseline to assess the improvements of our Track-CNN approach. Cells tracked by either Track-CNN or our heuristic method were also curated by human researchers using syGlass virtual reality software (Pidhorskyi et al., 2018) to assess the accuracy of tracking. Some of these curated traces were also returned to the training pipeline to improve our deep learning approaches in a positive-feedback loop (Figure 2A).

### Training data generation

All training data was curated by a human expert using syGlass software to provide point coordinates. To obtain volumetric segmentations, we trained an *ilastik* random forest regressor (Berg et al., 2019) to procure an over-sensitive voxel-wise segmentation model. Then, we excluded every *ilastik* identified object that did not overlap with a ground truth point coordinate to eliminate false positives in our over-sensitive *ilastik* model. Datasets were pooled from 12 animals and multiple treatment conditions. Image scales were standardized to 0.83 μm/pixel in XY and 3 μm/pixel in Z. Data was cropped to the appropriate input size for each respective neural network: Track-CNN 256 × 256 × 64 voxels, and seg-CNN 128 × 128 × 32 voxels.

Overall, Seg-CNN was trained with 6,828 training volumes and 759 validation volumes. Track-CNN was trained with 38,696 volumes and a validation set containing 4,300 volumes.

### Segmentation CNN training and inference

Seg-CNN employed a UNet architecture (Ronneberger et al., 2015) with 3D convolutional kernels built in Pytorch 1.6 (Paszke et al., 2017). The neural network took as input a 256 × 256 × 64 voxel volume containing fluorescently labelled oligodendrocytes in a single image channel (Figure 2B). The downsampling branch of the CNN contained 5 convolutional blocks with 5 × 5 × 5 filters, batch normalization, and max pooling to downsample the data and extract local features. The upsampling branch employed the same blocks in reverse. Max pooling operations were replaced by trilinear upsampling and 1 × 1 × 1 convolutions to resize the image back to the same input size while extracting global spatial features (Supplementary Figure S1). A final 1 × 1 × 1 convolution reduced the output to a two-channel volume which was softmaxed with a threshold of 0.5 to two classes corresponding to background and cell soma. Training was performed using a batch size of 2 for 30 epochs on an RTX 2080 Ti GPU. Loss was calculated as cross entropy and optimized using an Adam optimizer with weight decay (Loshchilov and Hutter, 2019) set at a learning rate of 10^−5^. During inference on unseen data, entire timeseries were fed to the neural network one timepoint at a time. Our algorithm then acquired 256 × 256 × 64 voxel crops from these volumes with 50% overlap to ensure all regions were assessed. Each crop was fed to Seg-CNN individually. The output segmentations of individual crops, with 50% overlap, were summed together and binarized before being stitched back into a full volume. The final analyzed timeseries is saved and returned to the user (Figure 2B).

### Track-CNN training and inference

Track-CNN employed a similar architecture to Seg-CNN except for a filter size of 7 × 7 × 7 for each convolution and a three channel 128 × 128 × 32 voxel input for our “seed-based” training approach. Seed-based training was employed to draw the attention of our CNN to individual cells in a volume by marking a cell of interest with a binary mask, or “seed” (Figure 3A). The input is thus a three-channel volume where channel 1 contains a raw fluorescence volume cropped from timepoint *t* and centered around a cell soma of interest. Channel 2 contains the binary mask/seed to indicate the cell of interest on timepoint *t*. All adjacent cells excluding the seed are set to a lower value. Finally, channel 3 contains a raw fluorescence volume cropped from timepoint *t + 1* but centered around the same position as in channel 1 (Figure 3A). In summary, this input provides the raw fluorescence from two consecutive timepoints and also indicates which cell we wish to track from timepoint *t* to timepoint *t + 1* using the binary mask in channel 2. Thus, the ground truth for optimization is a binary volumetric mask indicating the location of the cell of interest on timepoint *t + 1* (Figure 3A). Training was performed using a batch size of 4 for 18 epochs on an RTX 2080 Ti GPU. Loss was calculated as cross entropy and optimized using Adam optimizer with weight decay (Loshchilov and Hutter, 2019) set at a learning rate of 10^−5^ that was dropped to 10^−6^ at 13 epochs. During inference, volumes were cropped around each cell of interest in timepoint *t* along with seed masks and crops from timepoint *t + 1* to form a three-channel input for Track-CNN. This is repeated until all cells on timepoint *t* are assessed. Unassociated cells on *t + 1* are then added as newly formed oligodendrocytes to our list of candidate cells, and the analysis continues until all consecutive timepoints are tested (Figure 3A).

### Post-processing

To prevent misalignment of tracks, we included one major post-processing step in our analytic pipeline. We first noticed that, given a human tracked dataset, we could predict the location of a cell body on a subsequent timepoint within 0~10 pixels error by using the local directional vector of the tracks of five nearest neighbor cells from timepoint *t* to *t + 1* (Figure 3B, C). Thus, given that Track-CNN accurately tracks the majority of cells between consecutive timepoints, we can use the average local vector shift of the five nearest neighbors of any cell to correct for tracks that have severely gone off-target (> 12 pixel difference from predicted directional vector endpoint). These gross errors can then be re-evaluated. If an unassociated cell exists at the location of the predicted vector endpoint on *t + 1*, then the wrongly associated track now points to this unassociated cell. Otherwise, the track is terminated. We also included minor post-processing steps comprising of: (1) a minimum size threshold of 100 voxels for objects to be considered a cell soma; (2) objects that only exist on a single frame (excluding the first and last frame) are dropped, as they were likely to be debris.

### Heuristic baseline method

Since no baseline methods exist for comparison, we developed an approach to assess the extent to which deep learning outperforms traditional imaging informatics methods. We developed a tracking program in MATLAB R2020a (Mathworks) where cells are cropped from timepoint *t* and assessed on a pair-wise basis to identify whether its’ nearest neighbors on *t + 1* correspond to the same cell at timepoint *t*. To determine this association, we employed a few simple heuristics and rules: (1) successful tracking required a structural similarity index (SSIM) greater than 0.2 between cropped volumes from different timepoints. SSIM is an indicator of similarity that considers structure, intensity, and contrast-based differences between images. We applied the assumption that if a cell exists at *t + 1*, the overall local environment should look rather similar at timepoint *t*, thus a correct association would have a moderate to high SSIM. (2) Similar to the post-processing used for Track-CNN, we estimated the average vector of all nearest neighbors to model local tissue movement in a cropped field of view from *t* to *t + 1*. This allowed us to evaluate if the current track from *t* to *t + 1* flows in the same direction as the local shift of neighboring tracked cells. If the proposed track does not align with the local shift of neighboring tracked cells, then the track is terminated.

### SNR calculation

Since there is no standard for defining signal-to-noise ratio (SNR) in fluorescence imaging (Zhu et al., 2012), we adapted a standard logarithmic signal-processing SNR equation for our usage:

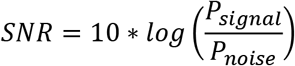

Where we defined *P*_signal_ as the average signal (meaningful input) and *P*_noise_ as the standard deviation of the background noise. However, since we have no reference image to define what perfect signal is in any raw dataset, we defined our signal to be any pixels above a certain value *j* and noise to be any pixels below that value.

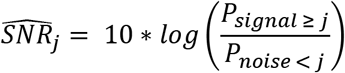

Where *P*_signal_ is defined as the mean of all values above *j*, and *P*_noise_ is defined as the standard deviation of all values below *j*. Since *j* would otherwise be arbitrarily determined, we chose to calculate *j* from the entire image volume using Otsu threshold for binarization (Otsu, 1979), providing us with a reference free metric of SNR.

### Statistical analysis

All statistical analysis was performed using Python statsmodels and scipy libraries. N represents the number of animals used in each experiment, unless otherwise noted. Data are reported as mean ± SEM or median ± SEM as indicated, and p < 0.05 was considered statistically significant. Level of significance is marked on figures as follows: * denotes p<0.05; ** denotes p<0.01; *** denotes p<0.001.

### Code availability

Packaged software code for *Oligo-Track* is readily available at github.com/Bergles-lab/Xu_Bergles_2021_Oligo_Track along with instructions for use. The algorithm is prepared to work independent of Linux and Windows operating systems, with minimum Python 3.6.

## RESULTS

### Quantifying oligodendrocyte dynamics *in vivo* using CNN-assisted cell tracking

To visualize individual oligodendrocytes in the cerebral cortex, cranial windows were surgically implanted in mice that express EGFP under control of the *Mobp* promoter/enhancer (Hughes et al., 2018; Orthmann-Murphy et al., 2020) (Figure 1A). Using two-photon microscopy, the somas and cytosolic processes of oligodendrocytes could be resolved up to a depth of ~400 μm from the pial surface (Figures 1B,C), providing the means to quantify changes in both the number and distribution of oligodendrocytes over weeks to months with repeated imaging. The dramatic increase in density of oligodendrocytes with depth (Figure 1C) presents challenges for unambiguous identification and increases the time necessary to mark and track cell positions throughout a time series. To overcome this quantitative challenge, we trained two sequential CNNs employing a UNet architecture (Supplementary Figure S1), which we termed *Seg-CNN* and *Track-CNN*, to follow oligodendrocytes *in vivo* during repetitive bouts of imaging over many weeks (Figure 2A).

**Figure 1:**
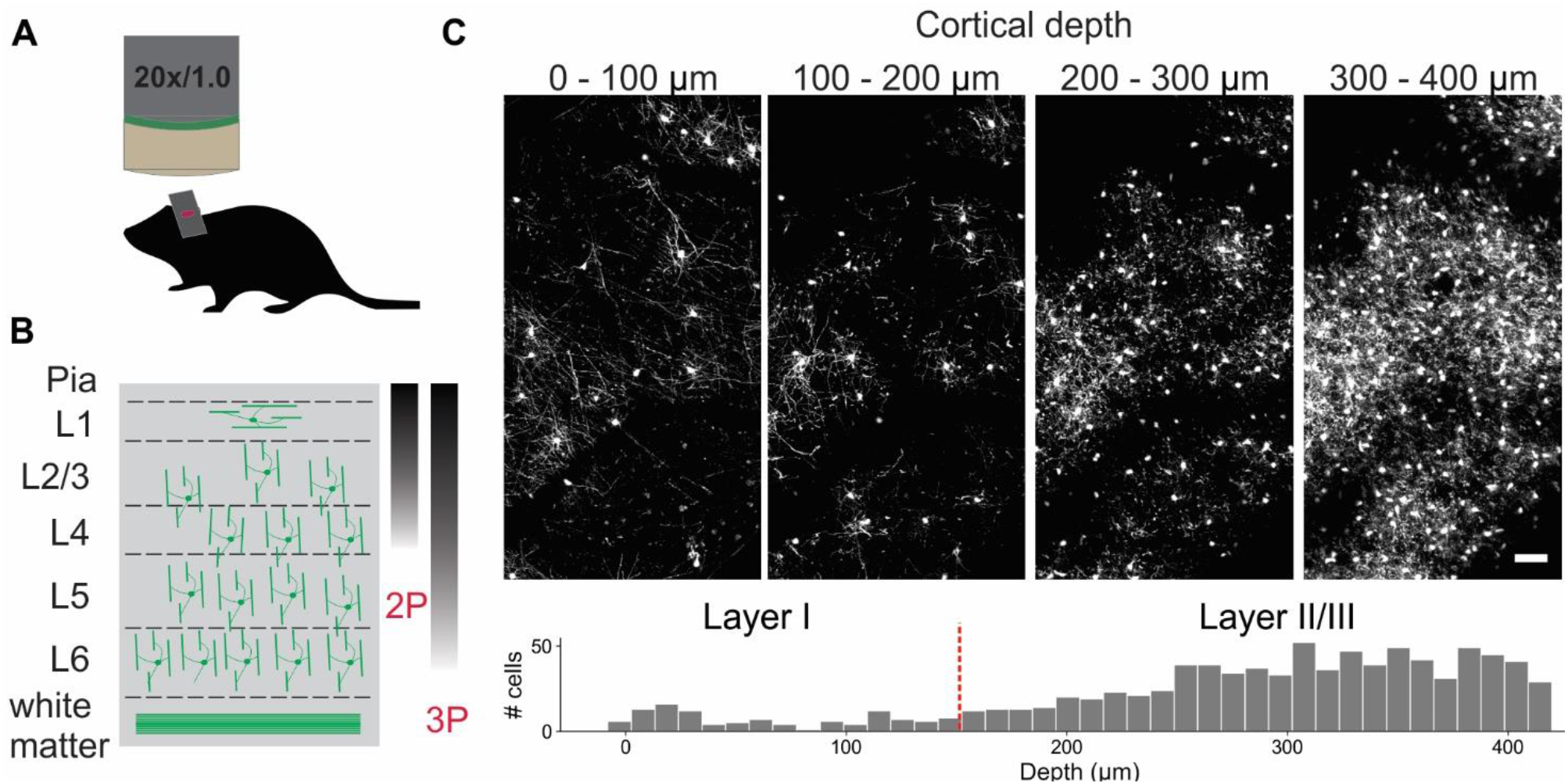
*In vivo* imaging of oligodendrocytes. (A) Cranial windows were surgically implanted in adult *Mobp-EGFP* mice in which only oligodendrocytes express EGFP. (B) Orientation of oligodendrocytes from imaging surface to white matter. Oligodendrocytes in upper cortical layers myelinate horizontally aligned axons, while those in deeper cortical layers are aligned perpendicularly to pial surface. Standard imaging range of two-photon and three-photon microscopy highlighted with approximate gradients (Theer and Denk, 2006; Lecoq et al., 2019) (C) XY maximum projections of 100 μm thick volumes at indicated depths (0 – 100 μm, 100 – 200 μm, 200 – 300 μm, 300 – 400 μm). Layer depths as estimated in somatosensory cortex (Narayanan et al., 2017). Oligodendrocyte density increases rapidly with depth, increasing the time needed for manual tracking. Scale bar: 50 μm.

**Figure 2:**
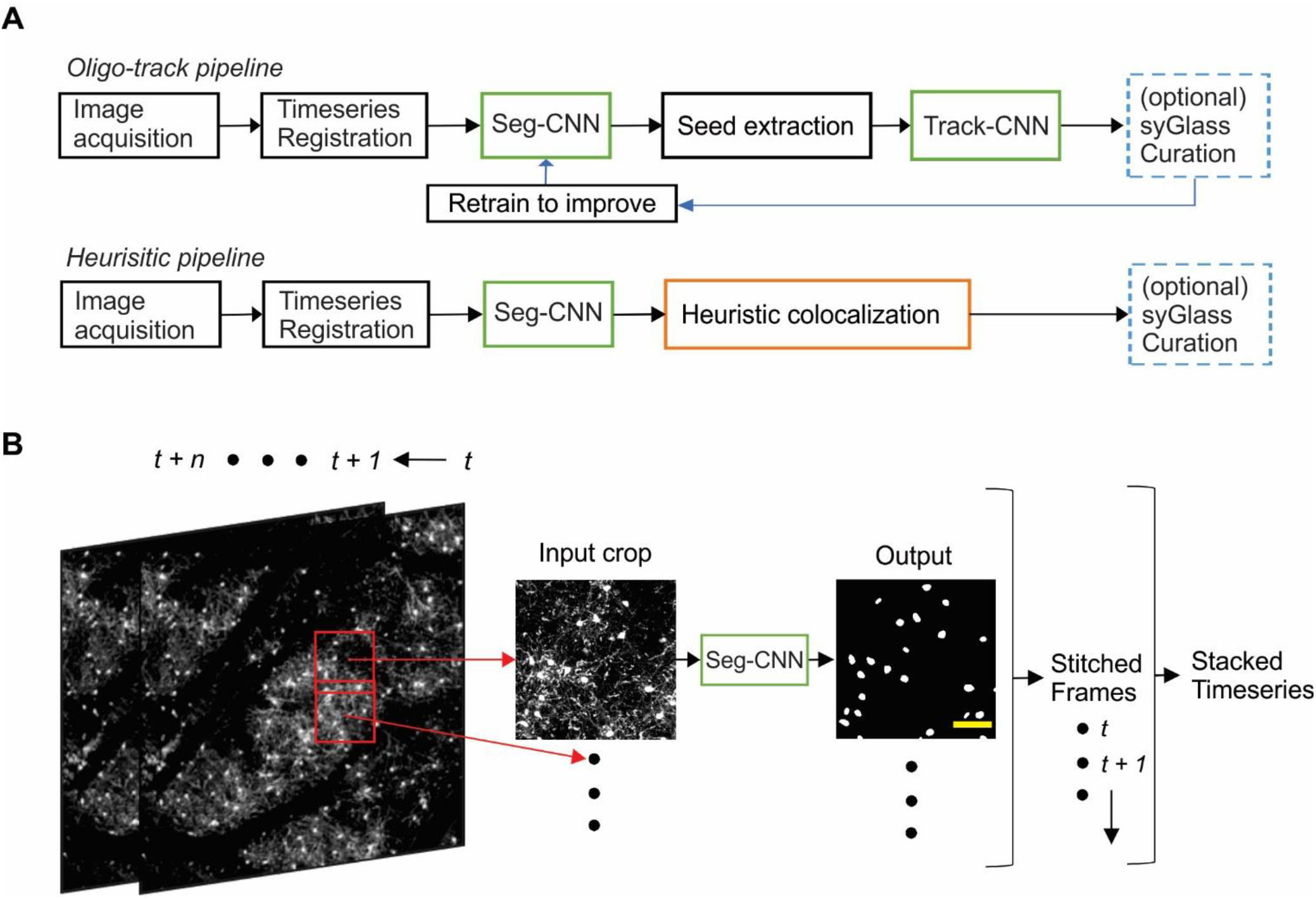
Computational neural network analysis pipeline. (A) Overview of the sequential CNN multi-object tracking pipeline *Oligo-Track* (top). CNNs marked in green. Overview of heuristic baseline method (orange) for comparison to *Oligo-Track* (bottom). Compatibility with optional syGlass curation provides validation of tracking in both pipelines (blue). Curated tracks can also be reintroduced into training pipeline for refinement of CNNs. (B) Seg-CNN pre-processing extracts cropped regions from larger volumes with 50% overlap for computational efficiency at each timepoint *t* to *t + n*. Cropped regions are restitched to form timeseries. Scale bar: 50 μm.

Images were first acquired over a 850 μm × 850 μm × 550 μm volume and then registered across time using ImageJ’s correct 3D drift plugin (Schindelin et al., 2012; Parslow et al., 2014) to adjust for small offsets. Seg-CNN was then used to perform semantic segmentation to identify the position of all oligodendrocyte cell bodies within the imaging volume at each timepoint in the timeseries. This process was completed sequentially on 256 × 256 × 64 voxel volumes that were adaptively cropped with 50% spatial overlap to reduce the amount of computer memory required to perform the computations (Figure 2B). The resulting binary segmentations were then re-stitched to create a stacked timeseries. Image stacks from sequential time points were then analyzed using Track-CNN, which employs a “seed-based” inference approach to determine whether any specific cell of interest exists in a subsequent timepoint. For all comparisons, we defined a tracked cell (or cell track), as a set of locations where a binary object was determined to be the same cell over subsequent timepoints by an algorithm or human researcher. The displacement vector for any cell thus starts at a soma on timepoint *t* and ends at the same tracked soma on *t + n*. Cell identification in Track-CNN is accomplished by providing a three-channel input to the CNN, which includes (1) a crop of raw fluorescence from timepoint *t* centered around a cell of interest, (2) a binary seed-mask that emphasizes the current cell of interest, and (3) a crop of raw fluorescence from timepoint *t + 1* that is centered around the cell on *t*. This allows the CNN output to be a volumetric segmentation of the same cell on timepoint *t + 1*, given a masked cell of interest on timepoint *t* (Figure 3A). Additional post-processing was performed using local tissue movement vectors to detect gross errors in tracking between sequential timepoints (Figures 3B, C). This post-processing used the observation that the displacement vector for any cell can be predicted within 10-pixel accuracy using the average displacement vectors of the nearest five tracked cells (Figure 3C). Thus, any cells with displacement vectors that varied drastically from predicted vectors, calculated from nearest neighbor tracks, could be classified as incorrect associations. Overall, during training, Seg-CNN performance plateaued after ~30 epochs, demonstrating accurate segmentation of cell somas relative to ground truth (Jaccard overlap index ~0.7) and detection of cells across all volumes (95% sensitivity, 91% precision; Supplementary Figure S2A). Track-CNN performance plateaued after ~5 epochs with highly accurate track associations (98% accuracy, 99% sensitivity, and 99% precision; Supplementary Figure S2B).

**Figure 3:**
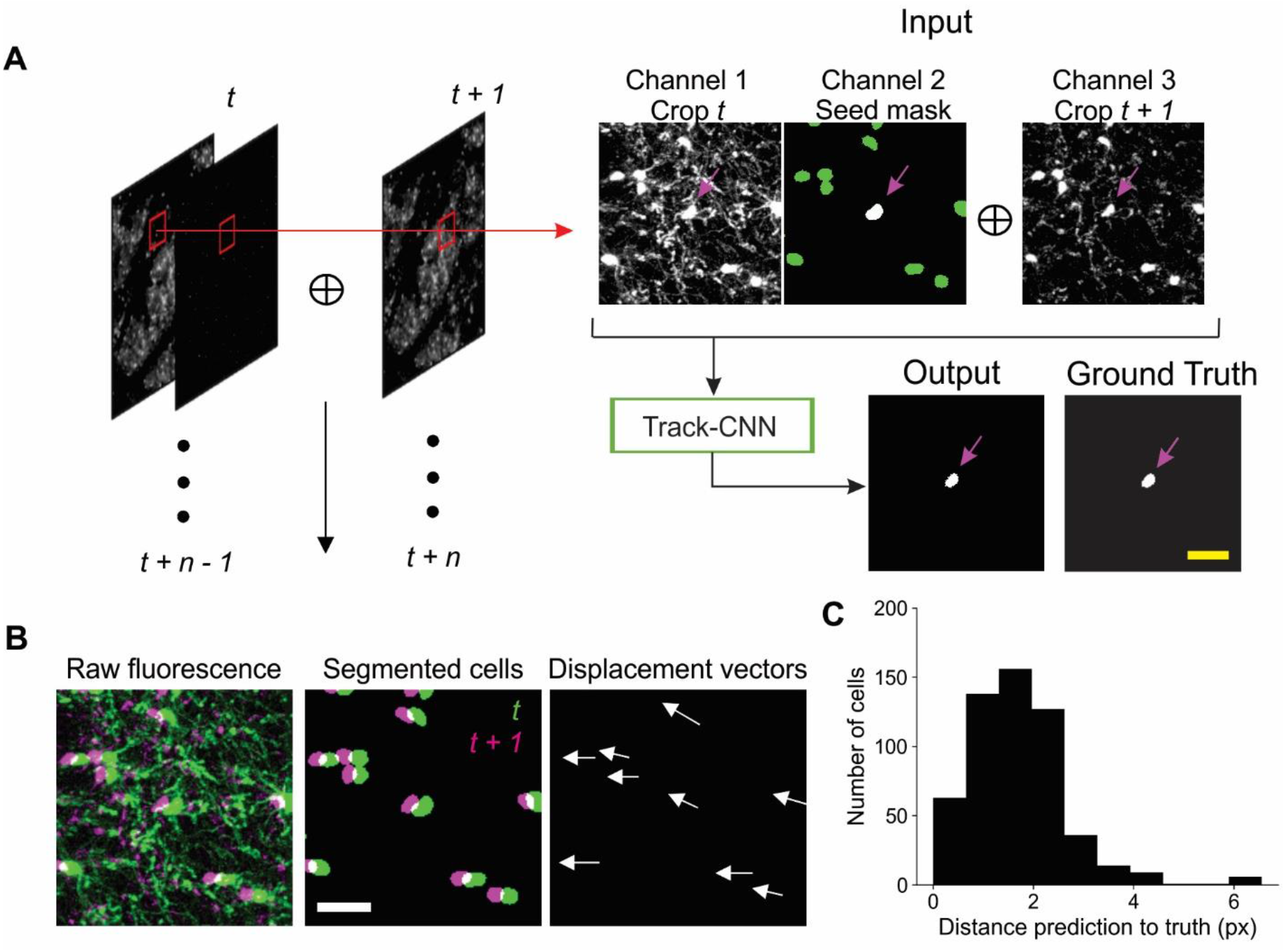
Track-CNN processing steps. (A) Crops are taken from each pair of timepoints *t* and *t + 1* centered around a cell denoted by magenta arrow on *channel 1*. *Channel 1* contains raw fluorescence from timepoint *t*. *Channel 2* contains seed mask of cell of interest (magenta arrow). Adjacent segmented cells are set to a lower value (green). *Channel 3* contains raw fluorescence from timepoint *t + 1*. Cropped images are concatenated together to form input to network. The network output is a semantic segmentation indicating the location of the seed masked cell on timepoint *t + 1*. This procedure is repeated for all cells on all consecutive timepoints. Scale bar: 30 μm. (B) Example showing local coherence in how tracked cells in a local region shift between timepoint *t* (green) and *t + 1* (magenta), allowing for predictive post-processing using average movement vectors (right). Scale bar: 30 μm. (C) Distribution of distances from predicted to actual location of cell on timepoint *t + 1* given any cell on timepoint *t*. The prediction is generated by taking the average displacement vector of five nearest neighbor tracks. Differences between predicted and actual location were typically within 6 pixels.

To determine if this CNN-based method outperforms a heuristic cell tracking method that employs similarity metrics and local tissue movement modeling, similar to the post-processing mentioned above, we tested both algorithms for their ability to extract biological trends of spontaneous cell regeneration in the cuprizone model of demyelination (Chang et al., 2012; Baxi et al., 2017; Hughes et al., 2018; Orthmann-Murphy et al., 2020). In this model, mice are fed cuprizone for three weeks, resulting in loss of >95% of oligodendrocytes in the upper layers of cortex, which are progressively regenerated as the mice are returned to a normal diet (Figure 4A). Both CNN and heuristic models detected the general trend of cell loss during the first three weeks of cuprizone treatment and subsequent oligodendrogenesis during recovery, as assessed relative to human counting (Figure 4B). However, closer examination revealed that *Oligo-Track* provided a more accurate accounting of cell dynamics. In particular, the heuristic method greatly mis-identified existing cells as being newly formed (Figure 4C), suggesting disrupted tracking. This conclusion was further supported by the increased number of wrongly terminated cell tracks by the heuristic algorithm at each timepoint (Figure 4D), suggesting that the heuristic approach often failed to identify existing oligodendrocytes in subsequent time points. We also assessed the difference in track length (persistence of cells during the time series) between ground truth and machine outputs (Figure 4E). Positive values in this plot indicate under-tracking, where the machine failed to track a cell in subsequent timepoints, while negative values indicate over-tracking, where the machine tracked a cell onto additional timepoints despite cell elimination determined in the ground truth. This graph reveals that Track-CNN markedly reduced the total rate of over-tracked segments errors two-fold from the heuristic algorithm (Figure 4F). Moreover, the severe error rate (under or over-tracking for > 1 timepoint) decreased almost five-fold. Together, these findings indicate that *Oligo-Track* provides substantial benefits for following oligodendrocytes in longitudinal 3D imaging datasets.

**Figure 4:**
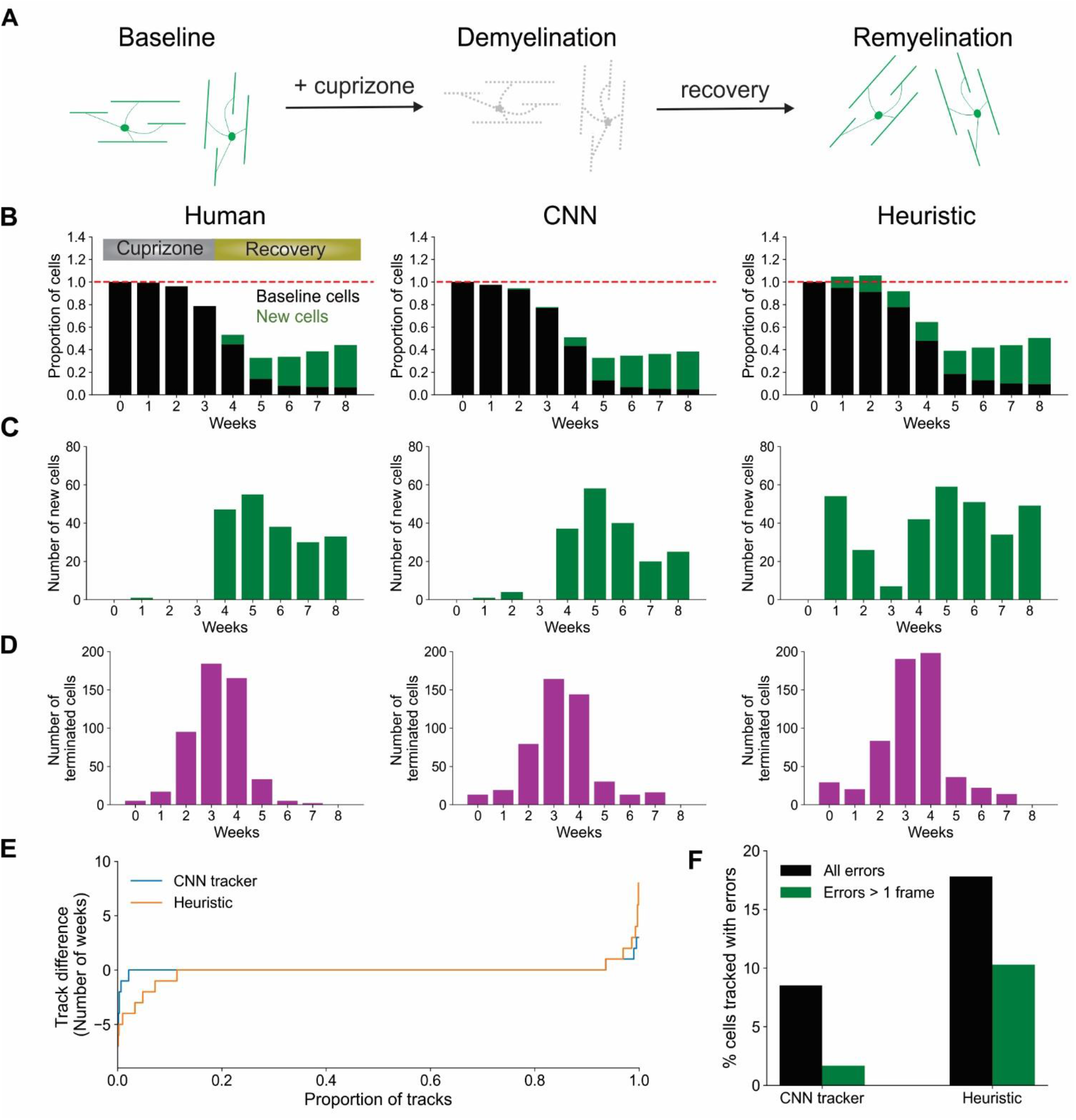
CNN-based tracking outperforms heuristic tracking. (A) Diagram illustrating cuprizone induced oligodendrocyte loss and recovery during the imaging period. (B) Overall normalized trends for human, CNN and heuristic tracking methods on test timeseries withheld from training data. (C) Number of new cells detected per timepoint for each method. (D) Number of cells terminated per timepoint for each method. (E) Track difference (length of track in ground truth - length of track by machine count) comparing ground truth to CNN and heuristic methods, respectively. (F) Comparison of major errors, defined as under- or over-tracking for > 1 timepoint, and total errors by CNN and heuristic methods.

### CNN-based analysis retains tracking ability despite changes in image quality

Many factors can influence image quality *in vivo*, limiting the ability to accurately assess cell dynamics. Cranial windows can become obscured by local inflammation at later (or earlier) timepoints, resulting in incorrect track associations by both humans and machines. Image scale, cellular debris, and laser power also commonly vary between experiments and impair implementation of standardized analyses. To assess the impact of these factors on our tracking algorithm, we started by first varying image scale, using bilinear interpolation to up- or down-sample raw data before performing Track-CNN analysis. The algorithm was mostly scale invariant, but struggled with up-sampling beyond two-fold (Figure 5A,B) showing that, optimally, input data should be scaled to the same 0.83 μm/pixel XY and 3 μm/pixel in Z resolution as the training dataset.

**Figure 5:**
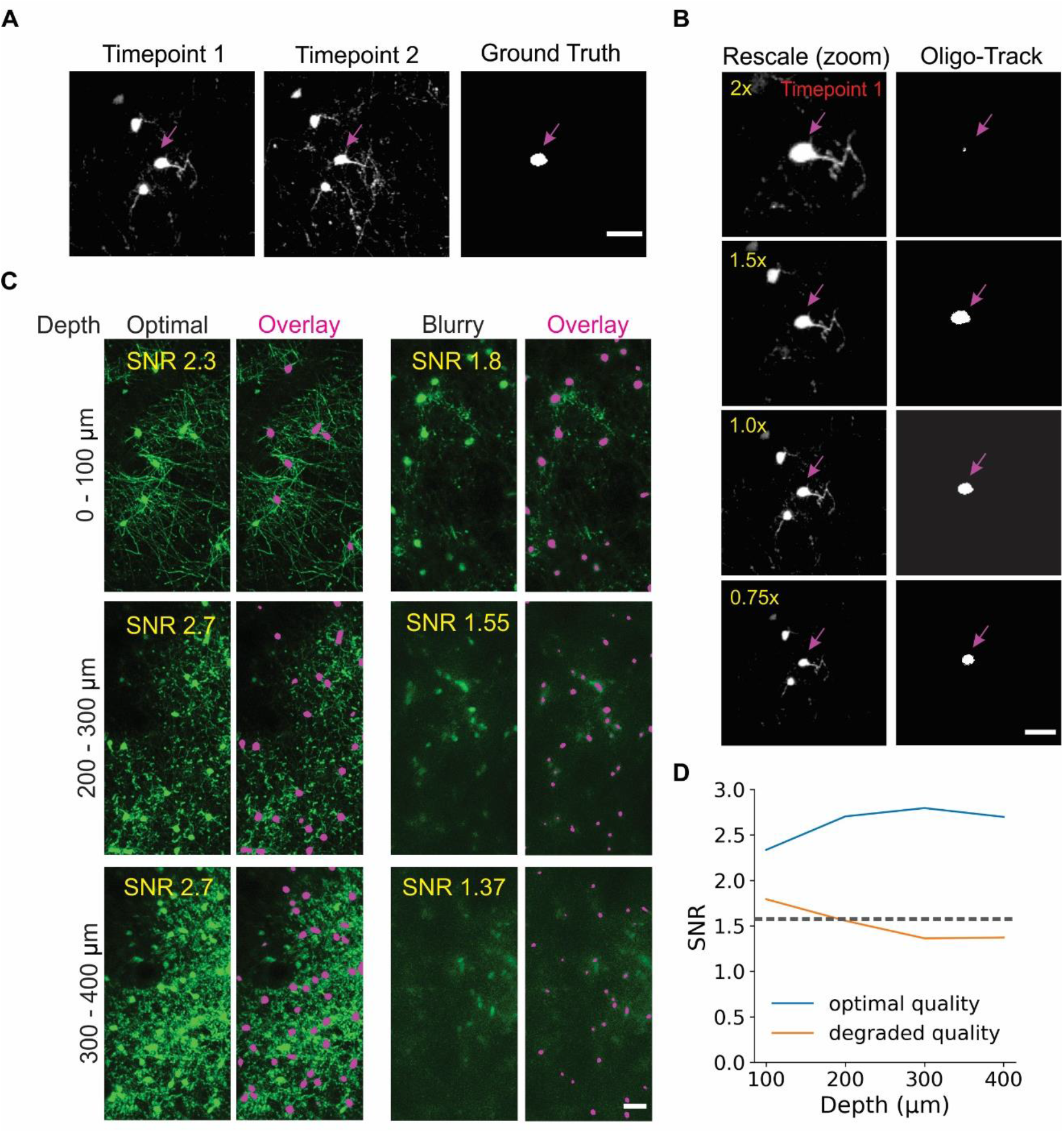
Oligo-Track enables robust cell tracking under different experimental conditions. (A) Input image for Track-CNN used in (B) to assess impact of different rescaling on Track-CNN performance. Cell of interest denoted by magenta arrow. Scale bar: 30 μm. (C) Representative XY maximum projections at indicated SNR values and depths in a volume with optimal image quality (left), and a volume with a less transparent cranial window (right). Overlay of cells detected by *Oligo-Track* in Magenta. Scale bar: 30 μm. (D) Plotting average SNR across depth of optimal quality and degraded quality volumes. Dashed grey line indicates human perceptual limit for reliably tracing data. Also represents point at which algorithm will provide warning to user.

We then assessed the impact of cranial window/image quality on tracking, using a custom reference free signal-to-noise (SNR) metric. We chose two representative imaging volumes, one from a mouse with an optimal cranial window, and one from a mouse with a window that had not yet become optically clear. The obscured window reduced the detection of fluorescence at lower cortical depths. Our average SNR metric clearly delineated the depth-dependent decay of image quality, as the SNR in maximum projections of the obscured volume dropped rapidly after a depth of 200 μm (Figure 5C,D). This image quality decay was verified visually, and while Seg-CNN still generalized and was able to identify oligodendrocyte somata in deeper layers despite the reduction in SNR, it was clear that many cells were obscured from view from both machine and human trackers (Figure 5C). By visual assessment, we set a threshold of SNR ~ 1.5 dB as a limit under which image quality becomes a concern for *Oligo-Track* analysis. Fluctuations in SNR between timepoints can lead to disrupted tracking as cells are arbitrarily obscured and falsely labelled as terminated or newly formed. This threshold was incorporated into our pipeline and offers users a warning during implementation of the algorithm.

Seg-CNN was also able to avoid some fluorescent, non-cellular components or weak cellular autofluorescence associated with cells other than oligodendrocytes, which can be difficult for non-deep learning approaches (Supplementary Figure 3A). However, the overwhelming density of brightly autofluorescent debris, such as lipofuscin found near the pial surface, were sometimes detected as false positives by Seg-CNN (Supplementary Figure 3B). We suggest that researchers using this software avoid areas with dense debris or lipofuscin, or at least exclude these regions from analysis, although this can be difficult when imaging in aged tissue (Moreno-García et al., 2018; A. Yakovleva et al., 2020). We also determined that while low imaging power impairs cell detection, post-hoc adjustments of the intensity histogram towards higher values recovered some undetected cells (Supplementary Figure 3C). Finally, we found that Track-CNN was robust to some variations in noise and motion blur. This was assessed by applying sequentially larger standard deviations of noise (10, 40, 50) and increasing the rotation range of random motion artifacts (4, 6, 10 degrees) using the *Torchio* python library (Pérez-García et al., 2021) (Supplementary Figure 3D,E). Together, this analysis shows that *Oligo-Track* can maintain performance despite changes in environmental variables that affect the distribution of the data. Moreover, we demonstrated that pre-processing of input data, such as intensity adjustments and the exclusion of regions with high debris or low SNR, can reduce instances of inaccurate tracking.

### CNN detects layer-specific suppression of oligodendrogenesis at extended depth

To assess the capacity of our pipeline to extract biological trends, we used the fully automated system to analyze oligodendrocyte dynamics for up to 12 weeks in cuprizone treated and non-treated control mice. As anticipated, cuprizone treatment resulted in a predictable time course of oligodendrocyte degeneration and subsequent regeneration after mice were no longer exposed to the drug, while control mice gradually added oligodendrocytes over several weeks (Orthmann-Murphy et al., 2020) (Figures 6A,B, Videos 1 and 2). Moreover, when cells were segregated into 100 μm thick blocks from the pial surface, greater suppression of oligodendrocyte regeneration was observed in the deeper layers of the cortex (Figure 6C), as reported previously (Orthmann-Murphy et al., 2020). The sensitive detection of *Oligo-Track* allowed rapid extension of the analysis by another 100 μm (300 – 400 μm block), revealing that regeneration was even less efficient than in the area above, providing further evidence of the depth dependent decline in oligodendrocyte regeneration in the somatosensory cortex.

**Figure 6:**
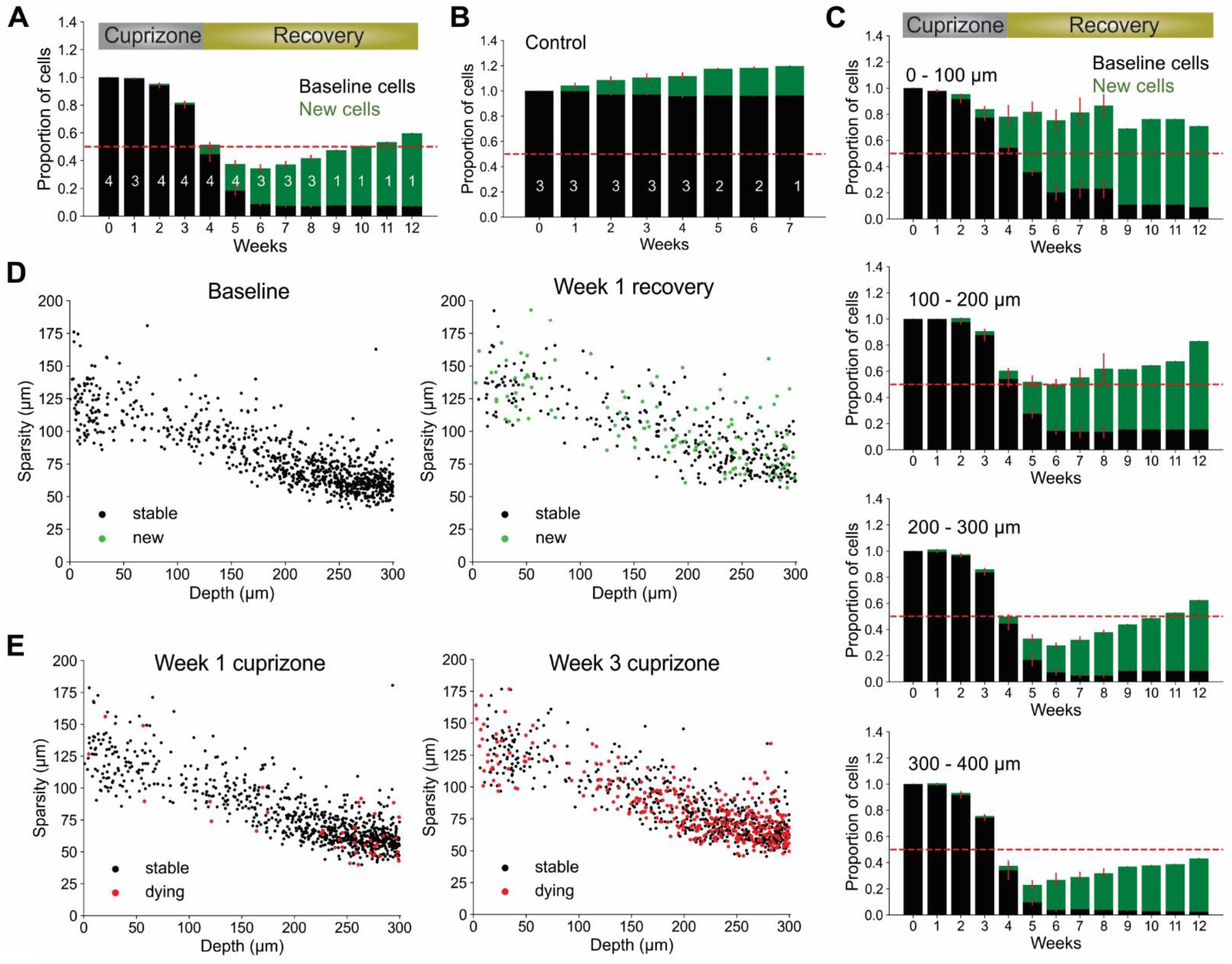
Local oligodendrocyte density does not correlate with region specific suppression of regeneration. (A) Normalized values for baseline and newly formed oligodendrocytes across weeks of cuprizone treatment and recovery. Bars indicate cell numbers averaged across animals for each week. (B) Normalized values for baseline and newly formed cells in no treatment condition. (C) Cortical depth-specific changes in oligodendrocyte regeneration showing suppressed regeneration in deeper layers. Volume split into 4 sections based on depth (0 – 100 μm, 100 – 200 μm, 200 – 300 μm, 300 – 400 μm). (D) Cell sparsity (average distance to 10 nearest neighbors) of stable and newly formed oligodendrocytes at baseline and week 2 of recovery shows no obvious clustering patterns at any timepoint. (E) Sparsity of cells that will die within one week (red) at week 1 and week 3 of cuprizone also shows no obvious clustering patterns at any timepoint. Cells pooled from n = 4 cuprizone treated and n = 3 control mice.

It is possible that the higher demand for oligodendrocyte regeneration in deeper cortical layers outstrips the regenerative capacity of OPCs (Hughes et al., 2013; Streichan et al., 2014). If the extent of oligodendrogenesis is limited by the availability of local cues or accumulation of myelin debris, then newly generated cells should preferentially appear in regions with lower initial oligodendrocyte density (and lower oligodendrocyte death) (Orthmann-Murphy et al., 2020). Our prior studies indicate that new oligodendrocytes do not regenerate in locations where previous cells had died, suggesting possible inhibition of proliferation by myelin debris after cell death (Lampron et al., 2015; Gruchot et al., 2019). As a measure of sparsity, we calculated the average distance from each cell to its five nearest neighbors. We limited our analysis to the first 300 μm of the cortex to avoid errors in sparsity calculations due to the lack of tracked nearest-neighbor cells past 400 μm depth. Given this measure, we found that there was no strong correlation between sparsity, cell death or regeneration (Figures 6D,E and Video 3), suggesting that cell death and regeneration are not strongly influenced by local oligodendrocyte density at baseline. Rather, global gradients of inhibitory factors such as cytokines released by astrocytes, which become persistently reactive in deeper layers of the cortex after cuprizone mediated demyelination (Orthmann-Murphy et al., 2020), may inhibit oligodendrocyte precursor cell differentiation (Skripuletz et al., 2008; Zhang et al., 2010; Su et al., 2011; Chang et al., 2012; Kirby et al., 2019).

### Volumetric segmentation enables identification of newly born oligodendrocytes

Oligodendrocytes undergo dramatic morphological changes as they transition from progenitors to mature myelinating cells, accompanied by an elaboration of myelin forming processes and changes in soma size (Kuhn et al., 2019). To quantify the time course of these somatic changes, we analyzed volumetric morphological data provided by *Oligo-Track*, from longitudinal imaging datasets where the birth date of newly formed oligodendrocytes was known. We limited our investigation to the first 300 μm of the cortex as tissue refraction often reduced brightness of cells in deeper cortical layers, resulting in inaccurate measurement of cell soma volume from dim fluorescence. This analysis revealed oligodendrocyte soma size was highly correlated with cell age. Most newly formed oligodendrocytes had larger cell bodies than stable cells at any timepoint across all depths (Figure 7A and Video 4). Projecting this across cell age, the soma volume of newly formed oligodendrocytes decayed exponentially over subsequent weeks from first appearance (Figure 7B,C; p < 0.001 @ 1 week, p=0.027 @ 2 weeks; Kruskal-Wallis test with Dunn’s post-hoc analysis). Moreover, the average volume of newly generated cells, post-cuprizone injury, was significantly higher compared to stable mature cells in control animals up to 3 weeks after oligodendrogenesis (Figure 7D; 1.6 ± 0.04 fold change p<0.001 and D=1.29 @ 1 week, 1.4 ± 0.04 fold change p<0.001 and D=0.85 @ 2 weeks, 1.2 ± 0.03 fold change p<0.001 and D=0.33 @ 3 weeks, 1.0 ± 0.03 fold change p=0.39 and D=0.07 @ 4 weeks; Kruskal-Wallis test with Dunn’s post-hoc analysis and Cohen’s effect size calculation). To confirm that this size difference is not associated with cuprizone induced changes, we also compared the volume of spontaneously generated oligodendrocytes in control animals with their stable counterparts and found that newly formed cells also had significantly larger cell somata (Figure 7D; 1.7 ± 0.09 fold change p< 0.001 and D=1.22 @ 1 week, 1.4 ± 0.07 fold change p<0.001 and D=1.1 @ 2 weeks, 1.3 ± 0.06 fold change p<0.001 and D=0.54 @ 3 weeks, 1.0 ± 0.05 fold change p=0.28 and D=0.09 @ 4 weeks; Kruskal-Wallis test with Dunn’s post-hoc analysis and Cohen’s effect size calculation). Thus, the increased soma size of newly formed oligodendrocytes is an innate biological phenomenon, rather than a response to cuprizone exposure.

**Figure 7:**
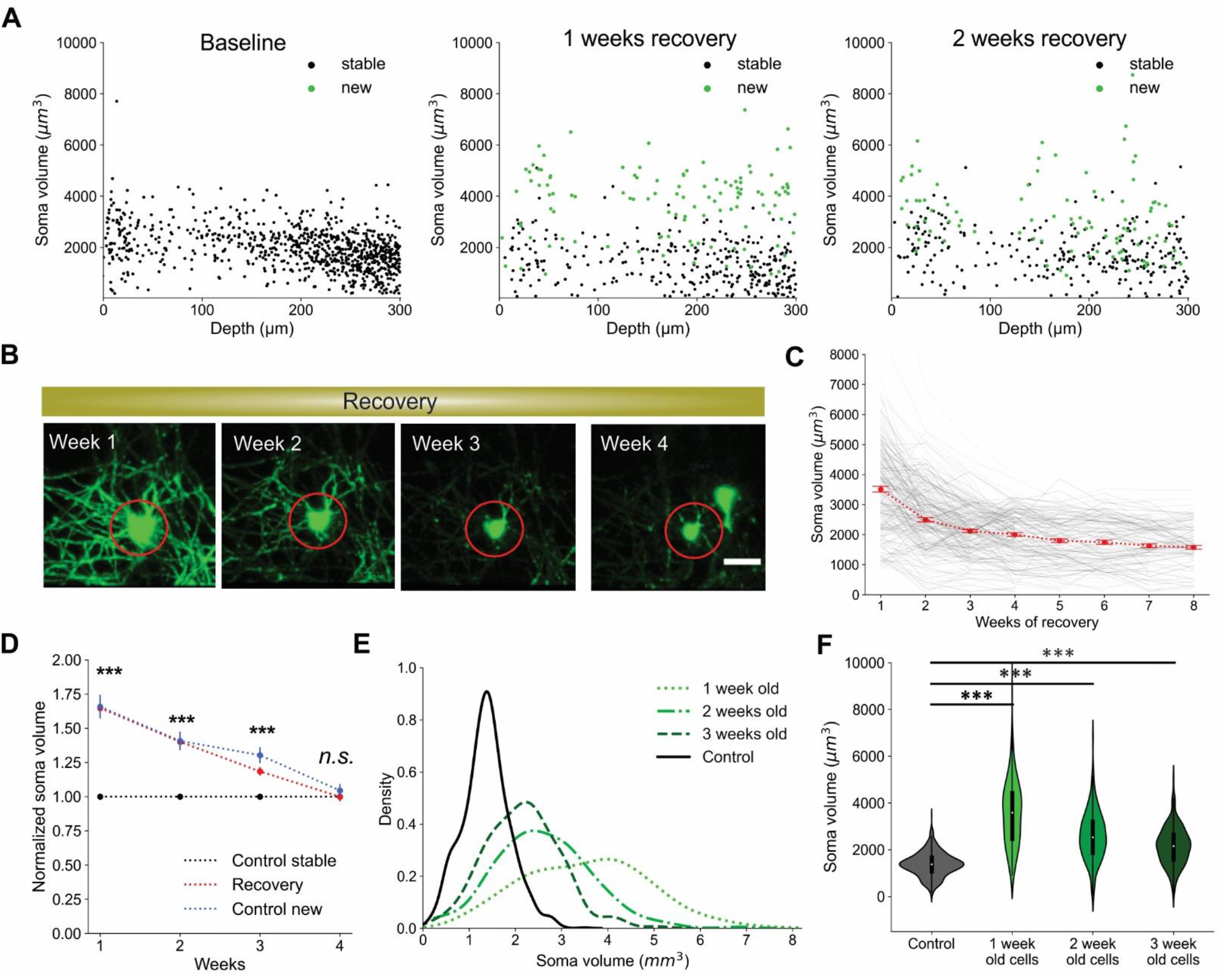
Newly generated oligodendrocytes can be identified by cell soma volume. (A) Soma volume of stable and newly formed oligodendrocytes at baseline, 1 week recovery, and 2 weeks recovery across different cortical depths. (B) Representative example of the change in soma size of newly formed oligodendrocyte tracked across 4 weeks of recovery. Scale bar: 20 μm. (C) Plot of decrease in soma volume for 250 cells over weeks relative to time of cell generation. Red dots are mean ± SEM. (D) Comparison of soma volume in newly formed oligodendrocytes during recovery compared with stable cells in non-treated mice. Also includes comparison of soma volume between spontaneously formed oligodendrocytes and stable cells in non-treated mice. All values are normalized to the mean soma volume of stable control cells at each matched timepoint. (E) Kernel density estimate for each distribution of soma volumes at indicated timepoints. These normalized distributions help visualize the probability that a cell with a certain soma volume is within a certain age range post-oligodendrogenesis. (F) Distribution of soma volumes of cells that are 1, 2, and 3 weeks old relative to cells in mice that are not treated with cuprizone at matched timepoints. Cells pooled from n = 4 cuprizone treated and n = 3 control mice. See Supplementary file 1 for statistical tests and significance level for each comparison.

Given the substantially larger cell somas of newly formed oligodendrocytes, we assessed the predictive power of cell soma size as an indicator of cell age. To examine the probability that a cell soma of a certain volume is exactly a certain age or within a range of ages, we plotted the kernel density estimate (KDE) for each distribution of soma volumes at different timepoints (Figure 7E). The KDE offers a normalized estimate of the probability density function such that we can visualize the probability of multiple conditions simultaneously. For example, we observed that a cell with a soma volume greater than 5000 μm^3^ has an almost 100% chance of being exactly 1 week old from time of differentiation. Similarly, cell somata between the range of 3500 - 5000 μm^3^ are most likely less than 2 weeks old, while somata larger than 3000 μm^3^ are likely newly generated cells within the first 3 weeks post-differentiation (Figure 7E). Finally, by comparing the mean soma volume of stable control oligodendrocytes to newly formed cells at multiple timepoints, we also confirmed the statistical significance of the predictive relationship between soma volume and cell age (Figure 7F; p<0.001 all comparisons; 1-way ANOVA with Tukey’s Honest Significant Difference post-hoc test).

### Oligodendrocyte death can be predicted from soma size

Oligodendrocyte death is typically preceded by nuclear condensation and shrinkage of the soma (Bortner and Cidlowski, 2002; Miller and Zachary, 2017). To determine if the soma size analysis could also be used to predict whether an oligodendrocyte will later degenerate, we plotted the soma volumes of all cells later observed to degenerate. After multiple weeks of cuprizone treatment, the median soma volume of all cells shrank significantly (Figures 8A–C, Video 4; p<0.001 @ 1 week, 2 weeks and 3 weeks), consistent with the high degree of oligodendrocyte degeneration observed in the cortex. When compared to oligodendrocytes at comparable timepoints in control mice, soma size was also significantly smaller after extended cuprizone treatment (Figure 8D; 0.83 ± 0.013 fold change p<0.001 and D=0.26 @ 1 week, 0.73 ± 0.013 fold change p<0.001 and D=0.65 @ 2 weeks, 0.6 ± 0.015 fold change p<0.001 and D=1.1 @ 3 weeks; Kruskal-Wallis test with Dunn’s post-hoc analysis and Cohen’s effect size calculation), consistent with progression to apoptosis. Given the large statistical power when sampling thousands of cells, we additionally defined a significant difference in soma volume as one having a medium to large effect size (> 0.5 Cohen’s D), which only occurred at 2 and 3 weeks of cuprizone treatment. Assessing the predictive power of soma volume again, we attempted to predict the likelihood that a cell would die within the next subsequent week given that the cell is smaller than a certain soma volume. While not as striking as the predictive power for newly formed oligodendrocytes, the probability that cells with somata below 500 μm^3^ would disappear within one week was over 90% (Figures 8E,F). Together, this analysis reveals that the size of oligodendrocyte somata calculated using deep neural networks can be used to predict, without prior or later longitudinal imaging data, whether a cell was recently generated and whether it is likely to degenerate.

**Figure 8:**
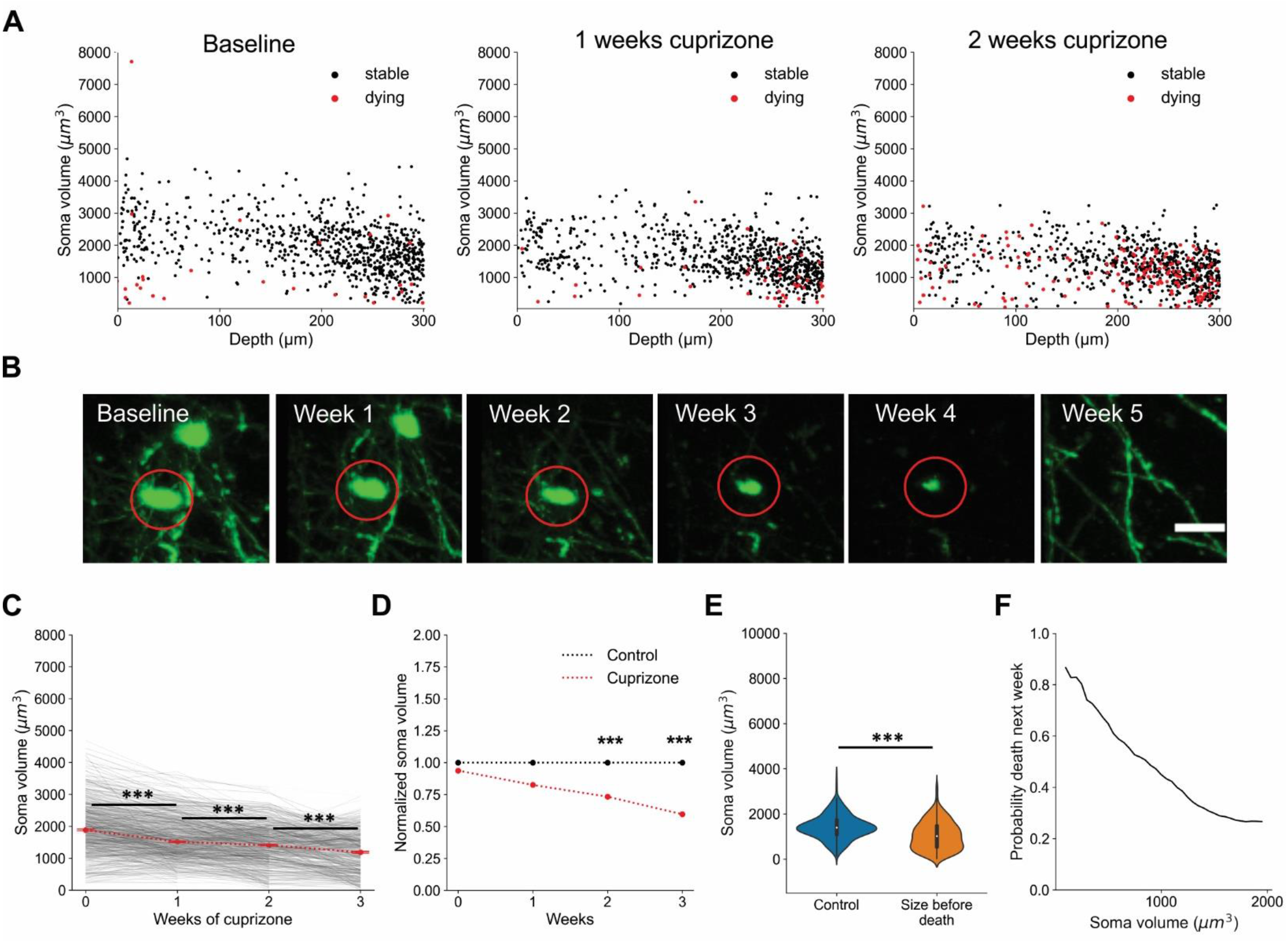
Oligodendrocyte death can be predicted from cell soma volume. (A) Volume of cell somata within 1 week of dying (red) at baseline, 1 week cuprizone, and 2 week cuprizone timepoints. (B) Representative example of cell soma shrinkage throughout cuprizone treatment, resulting in eventual death. Scale bar: 20 μm. (C) Plot of soma volume decrease for 860 cells during cuprizone treatment. (D) Plot of average soma volume of dying cells at each timepoint of cuprizone treatment relative to timepoint matched cells from control mice. All values are normalized to the mean soma volume of stable control cells at each matched timepoint. (E) Overall distribution of soma volumes for non-treated cells and cells within 1 week of death during cuprizone treatment. (F) Probability that a cell soma below a certain volume is within 1 week of death. Cells pooled from n = 4 cuprizone treated and n = 3 control mice. See Supplementary file 1 for statistical tests and significance level for each comparison.

## DISCUSSION

To facilitate analysis of oligodendrocyte dynamics in the adult brain we designed *Oligo-Track*, a deep learning pipeline that uses two sequential CNNs to allow cell tracking in volumetric imaging datasets. This methodology provides a substantial improvement over traditional imaging informatics approaches as it was faster, less subject to user bias and less influenced by factors that commonly deteriorate image quality, allowing reliable automated cell tracking over time series spanning multiple weeks. This automated volumetric analysis enabled us to increase the number of oligodendrocytes analyzed in deeper layers of the mouse cortex and to identify newly formed oligodendrocytes and those that are in the process of degenerating simply based on soma size at a single time point without longitudinal tracking information.

This CNN tracking pipeline follows a two-step approach to optimize multi-object tracking (MOT). We first setup a detection step, where oligodendrocytes are identified in a volume, followed by an association step, to link tracked cells across time frames (Ciaparrone et al., 2020). Unlike other deep learning MOT approaches, which often only use CNNs to generate bounding boxes or extract features (Ciaparrone et al., 2020), we employed two sequential CNNs that both performed semantic segmentation in the MOT detection and association stages (Seg-CNN and Track-CNN, respectively). The output of this pipeline provides not only the location of all tracked cells, but also the volume of each cell soma. This volumetric tracking was made possible by training our association network (Track-CNN) with a seed-based learning approach. Previous studies have shown that, when given input data containing several cells, one can mark cells of interest with a binary mask, or “seed”, to draw the attention of CNNs (Xu et al., 2019). This forces a semantic classifier to not only learn to identify oligodendrocyte somas, but also to identify the somas of individually marked cells of interest across different timepoints.

From a computational standpoint, there are several advantages to this automated approach. Roughly estimating the time for manual analysis with syGlass, a 3D virtual reality based visualization tool, we found that a 10-week, 10-timepoint dataset with a size of 800 × 800 × 300 μm per timepoint would take a researcher approximately six hours to identify and track all oligodendrocytes within this volume. This estimate only considers the time to place point coordinates and does not include the considerable additional time it would take to trace every voxel to generate volumetric segmentations. This estimate also does not consider how much longer manual analysis would take without access to specialized VR software (e.g. syGlass). By comparison, *Oligo-Track* requires ~20 minutes for Seg-CNN segmentation (~2 minutes per timepoint) and ~25 – 35 minutes for Track-CNN associations for the same volume across 10 timepoints, for a total analysis time of 45 – 55 minutes, more than six times faster than achieved with VR-assisted manual tracking, just for cell identification. This processing time is also purely computational, so manual labor time is reduced to almost zero, and offers fully volumetric segmentations. Total runtime will vary depending on cell density, number of timepoints, the size of volumes during inference and the exact computer configuration.

Standardization of methodology and accuracy are also important advantages of the CNN analysis approach. Losing dimensionality can be extremely detrimental to quantification speed and accuracy, as cells can often lie on top of one another or shift in unpredictable ways that can be missed if viewing 4D data in lower dimensional space. As many researchers do not yet have access to 4D visualization/tracking tools, *Oligo-Track* standardizes the approach to longitudinal cell tracking, removing the reliance on specialized proprietary software and reducing tracking inconsistencies between individuals.

Although there are clear technical advantages of using CNNs to track cells over time, the decision to use deep learning as an underlying analytic framework comes with additional considerations. Deep learning is often criticized for its “black box” nature, as researchers are unable to understand the intricate decision-making process of millions of weighted connections in a CNN, resulting in sometimes unpredictable behavior (Heaven, 2019; Yampolskiy, 2019).

For example, as we see in our own network, it was difficult to define the exact level of debris avoidance that the neural network was capable of, and why certain debris were more likely to be identified as false positives. This variability could be addressed in future work by data augmentation, whereby data containing high levels of real or synthetic debris could be introduced during CNN training. Currently, we partially addressed the unpredictability of deep learning by using VR-based 4D manual curation post-CNN analysis to ensure accuracy in unpredictable scenarios. We also used these post-hoc manually curated datasets to further improve the CNNs, highlighting a major advantage of deep learning approaches. CNN models are extraordinarily data hungry and can be continuously improved with new training data that help generalize to new imaging conditions (Klabjan and Zhu, 2020). For instance, while *Oligo-Track* has only been trained on cells up to 400 μm depth in the cortex, it will be possible to further train these networks to imaging conditions in deeper cortical layers. This training advantage is not available for traditional algorithms that may require extensive manual fine-tuning for extrapolation to slight variations in imaging conditions.

While the main limiting factor for developing deep learning technologies is the generation of large ground-truth training datasets to reach optimal performance levels, there are a growing number of methods by which researchers can reduce this high data demand of CNNs. For instance, transfer learning techniques have demonstrated how a network that is pretrained on a large dataset can be rapidly adapted to a new dataset with minimal new training data (Zhuang et al., 2020). Given the large database that our network was trained on, and the relatively similar features of cells that express fluorescent proteins, our pretrained CNN can serve as a basis for additional tool development, in which transfer learning is used to adapt this model to other cell types, where ground truth training data may not be readily available.

Automated quantitative tools will play a growing, critical role in the age of big data that is spurned by advances in biological imaging technologies. Of note for oligodendrocyte biology, three photon imaging promises to take us deeper *in vivo* (Horton et al., 2013; Lecoq et al., 2019), allowing us to examine the dynamics of these myelinating cells in layers 5 and 6 of the cortex and perhaps even into the white matter of the corpus callosum. Additionally, block-face imaging presents us with the opportunity to examine distributions of oligodendrocytes across the entire mouse brain, correlating myelination patterns with neuron type and brain region (Ragan et al., 2012; Amato et al., 2016; Winnubst et al., 2019). To match the scale of these imaging technologies, an important extension of the current work is to extract not only positional information about cells *in vivo*, but also the entire structure of cells. For oligodendrocytes, that means the soma, cytosolic branches, and individual myelin sheaths formed by each cell. As highlighted in this study, gaining quantitative access to even a single parameter, such as soma volume, can greatly extend biological understanding, allowing robust predictions to be made with limited data. Here, the strong correlations we observed between soma size, age, and survival provide us with a tool to infer the regenerative capacity of oligodendrocytes on fixed timepoint experiments acquired from individual tissue sections or from block-face imaging (Ragan et al., 2012). By extension, having access to the complete morphological structures of thousands of oligodendrocytes in the brain would enable us to assess complex region-specific differences in adaptive myelination, regenerative capacity and survival across the brain in mice subjected to different interventions.

Deep learning is well situated to provide us with the adaptable, reliable tools needed for the analysis of enormous new imaging datasets that can no longer be practically annotated using a manual brute force selection approach. Computational power is growing rapidly each year with new GPUs and the development of dozens of new deep learning techniques. Here, we demonstrate one powerful application of deep learning to resolve a multi-dimensional tracking challenge, which not only facilitates analysis of oligodendrocyte dynamics, but also extends our quantitative limits to extract novel insight into regional differences in regenerative capacity and allows predictions to be made about future behaviors. Having access to more cellular features and dynamics will bring us closer to understanding the events that underlie myelin regeneration that will aid in the discovery of therapeutics for treating demyelinating diseases.

## Supporting information

Supplementary Table of statistical analysis

## ACKNOWLEDGMENTS

We thank Dr. M. Pucak and N. Ye for technical assistance, T. Shelly for machining expertise, and members of the Bergles laboratory for discussions. Y.K.T.X. was supported by a fellowship from the Johns Hopkins University Kavli Neuroscience Discovery Institute and C.C. was supported by a National Science Foundation Graduate Research Fellowship. Funding was provided by NIH BRAIN Initiative grant R01 RF1MH121539, a Collaborative Research Center Grant from the National Multiple Sclerosis Society and the Dr. Miriam and Sheldon G Adelson Medical Research Foundation.

## Author contributions

Y.K.T.X. model design, conceptualization, data curation, formal analysis, supervision, validation, investigation, visualization, methodology, writing – original draft, writing – review and editing. C.C. conceptualization, data curation, supervision, methodology, writing – original draft, writing - review and editing. J.S. methodology, supervision, investigation, writing – original draft, writing-review and editing. D.E.B. conceptualization, Resources, Supervision, Funding acquisition, Investigation, Methodology, Writing – original draft, Project administration, Writing – review and editing.

---

**Video 1: Cell tracking across two stable timepoints.** Timepoint *t* (left) and *t + 1* (right). Magenta indicates the cell that is currently undergoing assessment by Track-CNN. After assessment, a color is assigned to the cell on *t* and *t + 1* to represent a tracked cell across timepoints. If the cell is untracked (or dies between timepoints), the cell soma is set to pure white on *t*. https://www.dropbox.com/s/nz9ll1n2ucw5v7w/Video_1_tracking_stable_cells_compressed.mp4?dl=0

**Video 2: Cell tracking across cuprizone injury timepoints.** Timepoint *t* (left) and *t + 1* (right). Magenta indicates the cell that is currently undergoing assessment by Track-CNN. After assessment, a color is assigned to the cell on *t* and *t + 1* to represent a tracked cell across timepoints. If the cell is untracked (or dies between timepoints), the cell soma is set to pure white on *t*. https://www.dropbox.com/s/zd647mqtnwsiokz/Video_2_tracking_cuprizone_cells_compressed.mp4?dl=0

**Video 3: Cell sparsity over weeks of cuprizone treatment and recovery.** Newly formed cells marked in green (left) and cells that will die within a week marked in red (right) starting from baseline followed by three weeks of cuprizone treatment and subsequent recovery. https://www.dropbox.com/s/vaem5qncd2jz5fh/Video_3_sparsity_over_weeks_compressed.avi?dl=0

**Video 4: Soma size of dying and newly formed cells over weeks of cuprizone treatment.** Newly formed cells marked in green (left) and cells that will die within a week a marked in red (right) starting from baseline followed by three weeks of cuprizone treatment and subsequent recovery. https://www.dropbox.com/s/vb2sgbilrcpuzdp/Video_4_volume_over_weeks_compressed.avi?dl=0

