## Supplementary Figures for "Automated in vivo tracking of cortical oligodendrocytes"

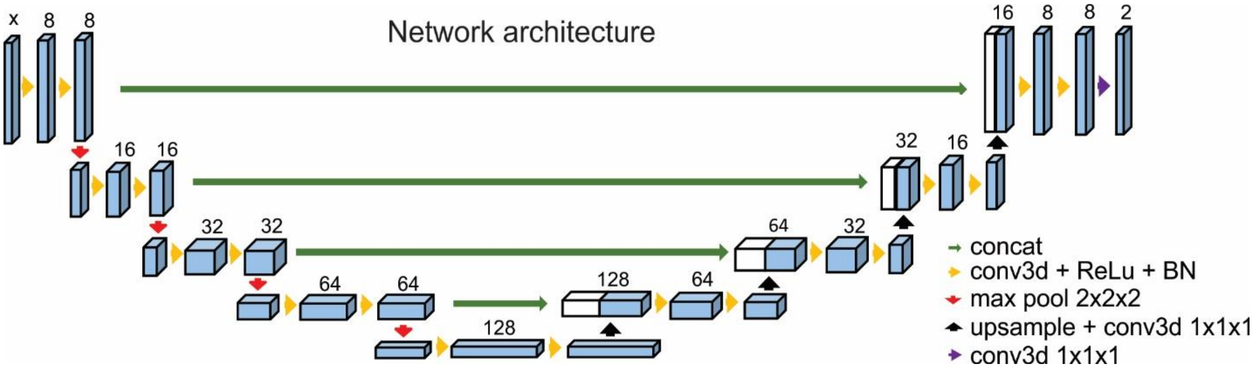


**Supplementary Fig S1: CNN architecture for Seg-CNN and Track-CNN.** UNet architecture used for both Seg- and Track-CNN. Downsampling branch extracts local features, upsampling branch extracts global spatial features.

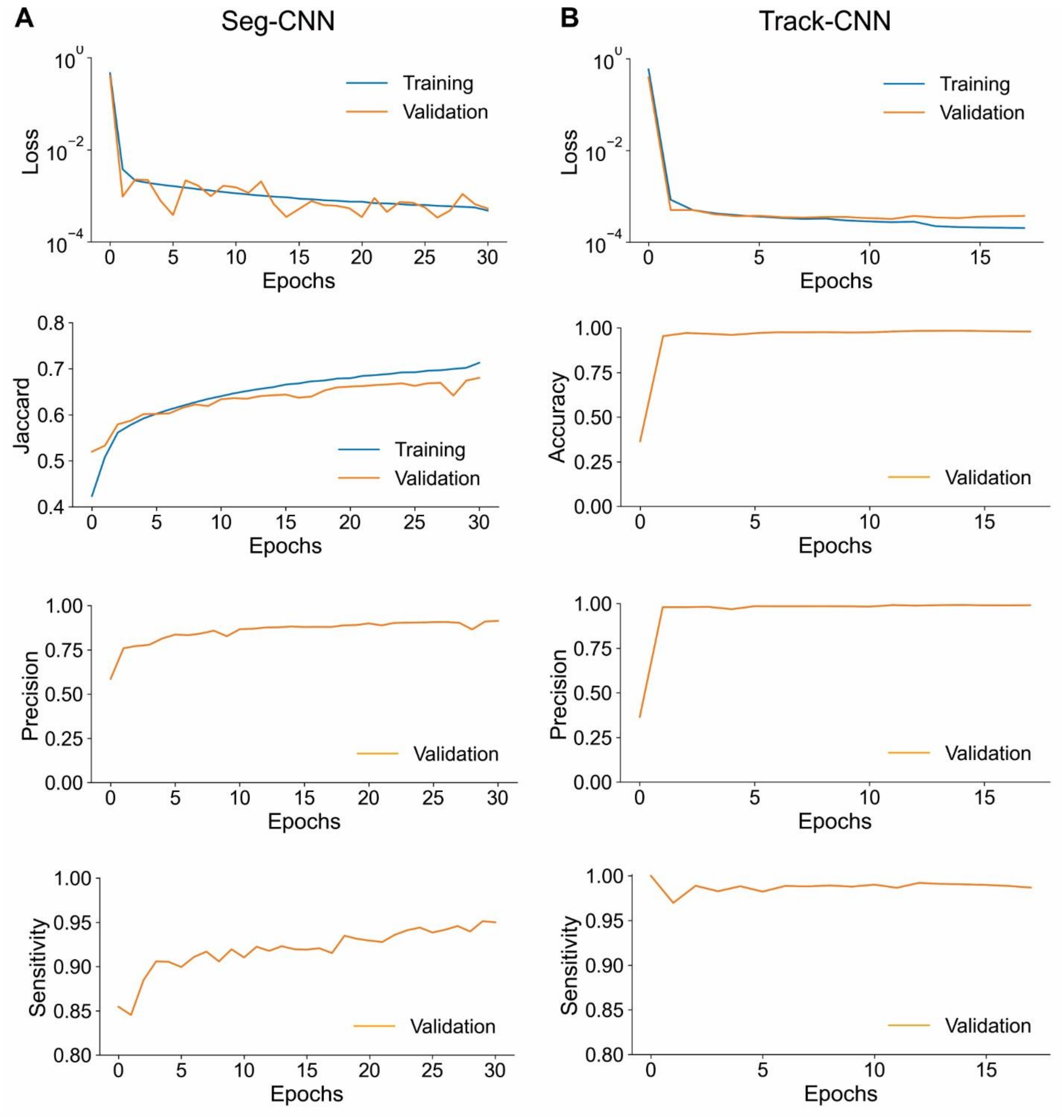


**Supplementary Fig S2: Training and validation performance for both CNNs.** (A) Seg-CNN loss, Jaccard overlap metric (Jaccard, 1912), precision, and sensitivity over 30 epochs. Curves associated with training and validation datasets are indicated. Loss descended stably without overfitting. All other metrics improved as anticipated. (B) Track-CNN loss, accuracy, precision, and sensitivity calculated per epoch. Accuracy, precision, and sensitivity only calculated on validation dataset for computational efficiency. Loss decreased stably without overfitting.

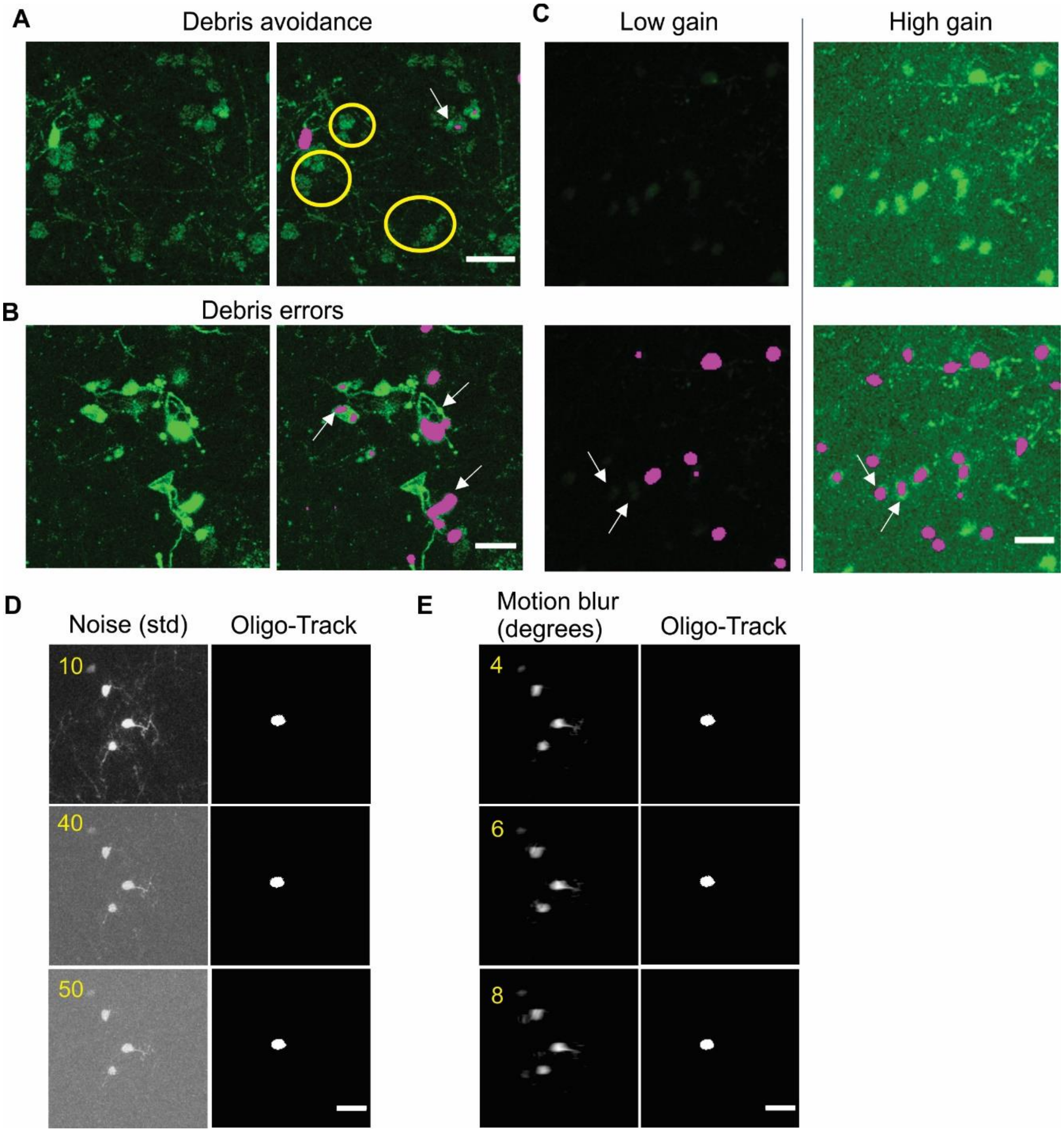


**Supplementary Fig S3: Robustness of CNN-based cellular tracking.** (A) Example of debris avoidance by Seg-CNN (yellow circles). Arrows indicate debris that was not avoided. Scale bar: 30 µm. (B) High intensity, dense debris is more likely to be detected as false positives. Arrows indicate debris that was not avoided. Scale bar: 30 µm. (C) Seg-CNN detections in image set with low laser power (left) and after gain adjustment (right). Arrows indicate initially undetected cells at low laser power. Scale bar: 25 µm. (D) Effect of different standard deviations of random noise on Track-CNN performance. (E) Effect of changing the rotation range (in degrees) of random motion artifacts on Track-CNN performance. Scale bar: 30 µm.
