## Supplementary Table of statistical analysis for "Automated in vivo tracking of cortical oligodendrocytes"

**Supplementary Table 1**: Summary of statistical tests and significance level for comparisons by comparison for each referenced figure panel.

| Figure panel | Comparison | Statistical test | Significance level |
|---|---|---|---|
| Figure 7C | Cell volume during recovery over weeks to first significant difference<br>@ 1 week vs. 2 weeks | Non-parametric Kruskal-Wallis H test with Dunn's post-hoc test | *p* = < 0.001 |
| | @ 2 weeks vs. 3 weeks | | *p* = 0.027 |
| | @ 3 weeks vs. 4 weeks | | *p* = 0.79 |
| | @ 4 weeks vs. 5 weeks | | *p* = 0.58 |
| | @ 5 weeks vs. 6 weeks | | *p* = 0.97 |
| | @ 6 weeks vs. 7 weeks | | *p* = 0.25 |
| | @ 7 weeks vs. 8 weeks | | *p* = 0.43 |
| Figure 7D | Cell volume of newly formed oligodendrocytes vs. stable control cells<br>@ 1 week | Non-parametric Kruskal-Wallis H test with Dunn's post-hoc test and Cohen's effect size (D) | *p* = < 0.001<br>*D* = 1.29 |
| | @ 2 weeks | | *p* = < 0.001<br>*D* = 0.85 |
| | @ 3 weeks | | *p* = < 0.001<br>*D* = 0.33 |
| | @ 4 weeks | | *p* = 0.39<br>*D* = 0.07 |
| | Cell volume of newly formed oligodendrocytes in control timeseries vs. stable control cells<br>@ 1 week | Non-parametric Kruskal-Wallis H test with Dunn's post-hoc test and Cohen's effect size (D) | *p* = < 0.001<br>*D* = 1.22 |
| | @ 2 weeks | | *p* = < 0.001<br>*D* = 1.10 |
| | @ 3 weeks | | *p* = < 0.001<br>*D* = 0.54 |
| | @ 4 weeks | | *p* = 0.28<br>*D* = 0.09 |
| Figure 7F | Mean distribution of control cell size vs. @ 1 week old cells | 1-way ANOVA with Tukey's Honest Significant Difference post-hoc test | *p* = < 0.001 |
| | @ 2 week old cells | | *p* = < 0.001 |
| | @ 3 week old cells | | *p* = < 0.001 |
| Figure 8C | Cell volume during cuprizone treatment over weeks<br>@ 0 week vs. 1 weeks | Non-parametric Kruskal-Wallis H test with Dunn's post-hoc test | *p* = < 0.001 |
| | @ 1 week vs. 2 weeks | | *p* = < 0.001 |
| | @ 2 week vs. 3 weeks | | *p* = < 0.001 |
| Figure 8D | Cell volume of cuprizone treated oligodendrocytes vs. stable control cells<br>@ 0 week | Non-parametric Kruskal-Wallis H test with Dunn's post-hoc | *p* = 0.001<br>*D* = 0.30 |

| | | | |
|---|---|---|---|
| | | test and Cohen's effect size (D) | |
| | @ 1 week | | *p* = < 0.001<br>*D* = 0.26 |
| | @ 2 weeks | | *p* = < 0.001<br>*D* = 0.65 |
| | @ 3 weeks | | *p* = < 0.001<br>*D* = 1.05 |
| Figure 8E | Mean distribution of control cell size vs. cells within 1 week of death | Unpaired 2-tailed t-test and Cohen's effect size (D) | *p* = < 0.001<br>*D* = 0.*65* |
